# More than stoichiometry: the molecular composition of inorganic and organic substrates controls ammonium regeneration by bacteria

**DOI:** 10.1101/2020.03.17.996322

**Authors:** J. Guo, M. Cherif

## Abstract

The mineralization of nitrogen (N) and especially the regeneration of ammonium are critical processes performed by bacteria in aquatic ecosystems. Quantifying these processes is complicated because bacteria simultaneously consume and produce ammonium. Here we use experimental data on the effects of the molecular composition of the supplied substrates, combined with a classical stoichiometric model of ammonium regeneration, to demonstrate how the quantification of these processes can be improved. We manipulated a batch culture experiment with an isolated bacterial community by adding three different types of N substrates: dissolved inorganic nitrogen (DIN, nitrate), dissolved organic nitrogen (DON, amino acid) and a mixture of DIN and DON. With such experiment set-up, the ammonium regeneration per se could be easily tracked without using complicated methods (e.g. isotope dilution). We compared the experimental data with the predictions of Goldman *et al*’ model (1987) as well as with a revised version, using the measured consumption carbon:nitrogen ratio (C:N ratio), rather than an estimated consumption ratio. We found that, for all substrates, and in particular, mixed substrates where C and N are partially dissociated between different molecules, estimates of ammonium regeneration rates can be improved by measuring the actual consumption C: N ratio.

**Importance:** Measuring bacterial ammonium regeneration in natural aquatic ecosystem is difficult because bacteria in the field simultaneously consume and produce ammonium. In our experimental design, we used nitrate as the inorganic nitrogen substrate. This way, we could measure separately the uptake and excretion of inorganic nitrogen by bacteria without incorporating cumbersome methods such as isotope dilution. Our experiment allowed us to evaluate the accuracy of various stoichiometric models for the estimation of net bacterial nitrogen regeneration. We found that:

1. The exact distribution of C and N among the various molecules that make the bulk of DOM is a crucial factor to consider for bacterial net nitrogen regeneration.
2. For all substrates, and in particular, mixed substrates where C and N are partially dissociated between different molecules, estimates of net nitrogen regeneration rates can be improved by measuring the actual C: N ratio of bacterial consumption.

## Introduction

Ammonium is a critical nitrogen (N) source for phytoplankton because this reduced substrate requires less energy to be assimilated than other sources of N such as nitrate (Glibert *et al.* 2016). In most ecosystems, ammonium results from mineralization by heterotrophic bacteria. They mineralize the N that is contained into the dissolved or particulate organic matter (DON and PON respectively) as ammonium and make it available for uptake by primary producers (Wheeler & Kirchman 1986). However, heterotrophic bacteria also take up ammonium or nitrate to build their own biomass. By taking up mineral N to meet their own needs, bacteria thus compete with primary producers (Tupas *et al.* 1994; Kirchman & Wheeler 1998; Glibert *et al.* 2016). The balance between bacterial ammonium regeneration and mineral N uptake, net N mineralization, thus determines the effect of bacteria on overall rates of primary production, and ultimately the productivity of higher trophic levels. Thus, it is a key ecosystem process (Danovaro 1998; Danger *et al.* 2007).

Positive net N mineralization usually occurs when the carbon and nitrogen ratio of the substrate used by bacteria (henceforth denoted *S*_*C:N*_) is lower than a given threshold C:N ratio, *T*_*N*_, the stoichiometric net mineralization threshold. Previous studies show that the threshold has a value around 10 for aquatic bacteria (e.g., net ammonium regeneration occurs when *S*_*C:N*_ < 10 : 1) (Parnas 1975; Goldman *et al.* 1987; Goldman & Dennett 1991; Cherif & Loreau 2007). When *S*_*C:N*_ > 10 : 1, then net immobilization of mineral N by bacteria may occur. These studies highlight the importance of *S*_*C:N*_ and *T*_*N*_ as a major factor determining whether bacteria take up or regenerate mineral N. However, what determines *T*_*N*_ is not fully addressed yet.

The net mineralization threshold obviously depends on *Y*_*C:N*_, the C: N ratio of newly-built bacterial biomass. In general, *Y*_*C:N*_ in aquatic bacteria is between 5:1 to 7:1 (mol:mol), which is lower than the net mineralization threshold (Cotner *et al.* 2010). Thus, other parameters besides *Y*_*C:N*_ must play a role in determining the threshold.

Mass-balance considerations dictate that:

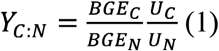

where *U*_*C*_ is the total amount of carbon consumed by bacteria, *U*_*N*_ is the total amount of nitrogen consumed, *BGE*_*C*_ and *BGE*_*N*_ are the bacterial growth efficiencies for C and N respectively, i.e., the fraction of consumption that is accumulated into biomass:

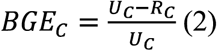

where *R*_*C*_ is the carbon respired, and

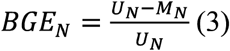

where *M*_*N*_ is ammonium regeneration by bacteria.

By combining (1) and (3), we can express ammonium regeneration *M*_*N*_ as a function of *Y*_*C:N*_, bacterial biomass C:N ratio, and bacterial growth efficiency for C, *BGE*_*C*_:

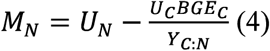

The equation (4) simply expresses the nitrogen mineralized, as the difference between the nitrogen consumed *U*_*N*_ and the nitrogen assimilated in proportion to the carbon accumulated in the biomass.

Measuring all components of the equation within the same experiment is rarely done in the field. In particular, because bacteria in the field simultaneously consume and produce ammonium, it is difficult to measure *M*_*N*_ and *U*_*N*_ at the same time without resorting to cumbersome methods based on isotope dilution (Tupas & Koike 1991). Therefore, it is usually assumed that the substrate is homogeneous and that the ratio of C to N uptake simply reflects the ratio of C-to-N in the substrate (Goldman *et al.* 1987):

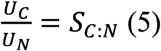

we define the net N mineralization threshold, *T*_*N*_ as

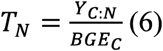

where *T*_*N*_ is approximately 10:1 in many aquatic ecosystems (Goldman *et al.* 1987; Goldman & Dennett 1991). Hence, under the assumption in (5) (6), (4) becomes (as expressed in Goldman *et al* 1987):

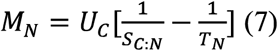

One can notice that the term between brackets in equation (7) can be positive or negative, corresponding to either immobilization or mineralization depending on whether *S*_*C:N*_ is smaller or larger than *T*_*N*_. Because ammonium is an easily available source of N to bacteria in most aquatic ecosystems (Kirchman 1994), it is frequently used as substrate in controlled experiments (such as in Goldman *et al* 1987). When ammonium is both a substrate and an excretion product, it becomes almost impossible to measure ammonium regeneration (*M*_*N*_) separately from nitrogen consumption (*U*_*N*_) and only the resultant net nitrogen mineralization (*NNM*) can be measured:

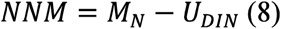

where *U*_*DIN*_ is the dissolved inorganic nitrogen consumed by bacteria.

Hence, equation (7) need to be modified in order to describe net nitrogen mineralization based on equation (8):

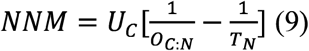

where *O*_*C:N*_ represents the C:N ratio of the organic fraction of the substrate. Here, it is usually assumed that the substrate is homogeneous and that the ratio of organic C to N uptake simply reflects the ratio of organic C to N in the substrate (Goldman and Dennett 1991):

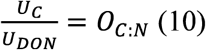

where *U*_*DON*_ is the organic N consumed by bacteria, often calculated as *U*_*DON*_ = *U*_*N*_ − *U*_*DIN*_.

Equation (9), expressed in slightly different forms, is the basis of most models that relate growth, substrate use and nutrient release in bacteria (e.g. Goldman & Dennett 2000; Daufresne & Loreau 2001; Manzoni *et al.* 2017).

Equation (9) has been tested successfully in the lab on bacteria that grow on labile organic substrates with similar bioavailability (Goldman *et al.* 1987; Goldman & Dennett 1991, 2000). The equation hinges on the simplifying assumption made in equation (10) that organic C and N are consumed in the same proportion as they are supplied. However, in natural waters, there is a wide diversity of C and N sources available for bacteria (e.g. amino acids, fatty acids) that differ widely in bioavailability, so that bacterial preference for different substrates could directly influence *U*_*C:N*._ Whether the C:N consumption ratio *U*_*C:N*_ completely reflects *O*_*C:N*_ (equation (10)) or not is thus questionable. An extreme case is when bacteria grow only on mineral N, i.e., the sources of C and N are totally dissociated. Then *U*_*C:N*_ is likely to reflect bacterial needs, and thus equal *T*_*N*_ rather than *O*_*C:N*_. Indeed, in cases where no N is supplied in organic form, i.e., *O*_*C:N*_ → +∞, equation (9) takes the simpler form:

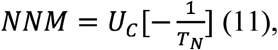

meaning that bacteria are predicted to immobilize nitrogen in proportion of their consumption of C and their relative biomass demand in C and N (*T*_*N*_).

Given the importance of equation (9), both empirically and theoretically, as the basis of bacterial *NNM* measurements used to quantify the stoichiometry of C and N cycles, we performed an experiment to assess its validity under different mixes of C and N resources. We grew an isolated bacterial community under 3 conditions 1) substrates where C and N are strongly coupled to each other, i.e., they are bound in the same DON molecule. For this situation we expect *U*_*C:N*_ to equal *O*_*C:N*_ and thus both equations (9) and (10) to hold; 2) substrates where C and N are perfectly dissociated, i.e. using nitrate and a DOC molecule as substrates. Here, we expect *U*_*C:N*_ to be decoupled from *O*_*C:N*_ and thus equation (11) to hold instead of equation (9); 3) substrates where C and N are partially associated, i.e., combining the first and second substrates. Here *U*_*C:N*_ is expected to reflect *O*_*C:N*_ only partially and thus equations (9) and (10) to apply only partially. The set-up of the experiment allows measuring *U*_*C*,_ *U*_*N*,_ *U*_*DIN*,_ *U*_*DON*_ and *M*_*N*_, so that consumption ratios can be calculated independently from the supply ratio *S*_*C:N*_. Hence, the experiment will enable us to conclude on the importance of using C and N consumption in models and assays of net N mineralization by bacteria, rather than using C and N availabilities as proxies.

## Materials and Methods

### Bacteria community isolation

We isolated a bacteria community from a non-axenic single strain phytoplankton (*Monoraphidium minutum*) batch culture. We assumed that a bacterial community is better equipped to use a variety of C and N resources than a single strain. The phytoplankton culture was grown in a Combo medium (Kilham *et al.* 1998) with only half of the nitrate concentration (500-µM-N). The culture was kept at 18°C under a 16:8 light: dark cycle for 2 weeks. The water was filtered through three GF/F glass fibre filters (0.7 µm pore size) to isolate the bacteria community from the phytoplankton and from other potential contaminating protozoa. After filtration, the absence of phytoplankton in the culture was checked by analyzing phytoplankton abundance in the culture with flow cytometry and the culture was immediately covered by foil and stored at 18°C for further manipulations.

### Experimental procedure

The bacteria isolated were inoculated (with a volume of 6 ml each) into bottles containing 600 ml of a COMBO medium (Kilham *et al.* 1998) modified so as not to contain nitrogen. N was added separately, using different substrates in different treatments: 1) DIN/DOC treatment: bottles received N as nitrate, ensuring that all nitrogen available to bacteria was in a dissolved inorganic nitrogen form (DIN); 2) DON treatment: bottles received an amino acid as the nitrogen source, ensuring that all nitrogen available to bacteria was in a dissolved organic form (DON); 3) COMBINED treatment: bottles received nitrogen in both inorganic (as nitrate) and organic (using the same amino acid) forms (DIN/DON). The final total N concentration in all the treatments was kept the same (100 µM-N). We chose a concentration of 100 µM-N to ensure that bacteria growth was not limited by mineral elements besides nitrogen. All the treatments received organic carbon as well, using different substrates in different treatments: 1) DIN/DOC treatment: bottles that received nitrate as their N substrate received an organic molecule containing C and no N; 2) DON treatment: bottles that received an amino acid as their N substrate already contained organic C in the amino acid, and did not receive further organic molecules; 3) COMBINED treatment: bottles that received both nitrate and an amino acid received the same organic compound as in the DIN/DOC treatment in order to complement the organic C already contained in the amino acid. The C-only containing molecule was carefully chosen so as to have chemical properties close to the amino acids selected, and take part in the same metabolic pathway within the bacteria. In order to test different substrate C:N ratios, we used two different amino acids: L-alanine and glutamate (Table 1). We chose pyruvate as the C-only substrate associated to alanine, and *α*-ketoglutarate as the C-only substrate associated to glutamate. All four molecules are substrates to the alanine-aminotransferase, an enzyme that is central to the metabolism of amino acids in most organisms (Mehta *et al.* 1993). The concentrations of C-only substrates added in the DIN/DOC and COMBINED treatments were calculated so as to yield the same concentration as in the corresponding DON treatment (Table 1). In total, we set 6 different treatments in the experiment, three with a C:N ratio of 3, containing combinations of nitrate, pyruvate and L-alanine, and three with a C:N ratio of 5, containing combinations of nitrate, *α*-ketoglutarate and glutamate (Table 1). Each treatment was replicated 4 times. Finally, we incubated 8 non-inoculated control bottles in parallel, containing COMBO medium, without any organic C substrate, in order to test for potential N losses, contaminations and other artefacts. The total number of bottles in the experiments was thus 32.

**Table 1.** Carbon and nitrogen substrates used in the different treatments. Each treatment was carried out with two different sets of molecules (nitrate/pyruvate/alanine, versus nitrate/*α*-ketoglutarate/glutamate) yielding two different total C concentrations (300 µM vs. 500 µM respectively) and two different C:N ratios (S _C:N_ =3 mol:mol vs. 5 mol:mol). N and C concentrations at the start of the experiment are indicated between parentheses.

| Total C:N<br>ratio | CN<br>molecular<br>composition | Treatments |  |  |
| --- | --- | --- | --- | --- |
|  |  | DIN/DOC | COMBINED | DON |
| $S_{\text{C:N}} = 3$ | N-only<br>and<br>C-only<br>substrates | Nitrate, $\text{NO}_3^-$<br>(100 $\mu\text{M-N}$)<br>Pyruvic acid, Pyr<br>(300 $\mu\text{M-C}$) | Nitrate, $\text{NO}_3^-$<br>(50 $\mu\text{M-N}$)<br>Pyruvic acid, Pyr<br>(150 $\mu\text{M-C}$) | |
| | Dual C-N<br>substrates | | L-alanine, Ala<br>(50 $\mu\text{M-N}$, 150 $\mu\text{M-C}$) | L-alanine, Ala<br>(100 $\mu\text{M-N}$, 300<br>$\mu\text{M-C}$) |
| $S_{\text{C:N}} = 5$ | N-only<br>and<br>C-only<br>substrates | Nitrate, $\text{NO}_3^-$<br>(100 $\mu\text{M-N}$)<br>α-ketoglutarate, Ket<br>(500 $\mu\text{M-C}$) | Nitrate, $\text{NO}_3^-$<br>(50 $\mu\text{M-N}$)<br>α-ketoglutarate, Ket<br>(250 $\mu\text{M-C}$) | |
| | Dual C-N<br>substrates | | Glutamate, Glu<br>(50 $\mu\text{M-N}$, 250 $\mu\text{M-C}$) | Glutamate, Glu<br>(100 $\mu\text{M-N}$, 500<br>$\mu\text{M-C}$) |

Bacteria were inoculated on day 0 and the bottles were closed by autoclaved cellulose stoppers that prevented bacterial contamination but enabled gaseous exchange. Cultures were maintained for 8 days at 18°C and under a16:8 light:dark cycle at 120 µmol photon m^-2^ s^-1^ light intensity. We sampled all bottles every second day to determine the abundance of bacteria. Chemical analyses were performed at the beginning and at the end of the experiment.

### Sampled variables

#### Bacterial abundance

Samples for bacterial abundance (BA) were taken at the start and end of the experiment, and every other day. 2 ml were preserved with 0.1% glutaraldehyde (final concentration) and immediately stored at −80 °C. The bacterial abundance was later measured with a FACSVerse™ flow cytometer (BD Biosciences) equipped with a 488 nm laser (20 mW output) and a 640 nm laser (output 40 mW). Frozen samples were quickly thawed in a 30°C water bath and stained with SYBR Green I (Invitrogen) to a final concentration of 1:10 000 (Marie *et al.* 2005). The samples were run at a flow rate of 40 μl min^-1^ during 1 min. When necessary, samples were diluted with sterile culture medium to avoid coincidence. Microspheres of 1 μm (Fluoresbrite plain YG, Polysciences) were added to the samples as internal standard. Forward light scatter (FSC), side light scatter (SSC) and green fluorescence from SYBR Green I (527 ±15) were used to discrimination bacteria.

#### Water chemistry and stoichiometry of bacteria

Total dissolved nitrogen (TDN), DOC, and DIN were determined at the beginning and at the end of the experiment. All the samples for the dissolved nutrients analyses were pre-filtered through 0.2 µm Filtropur syringe filters. For DOC and TDN analyses, samples were acidified with 1 ml of a 2M HCl solution. Then samples were analyzed by using an infrared gas analyzer (HACH IL-550 DOC-TDN). For DIN (NO_3_^-^ and NH_4_^+^), samples were analyzed with an automated flow injection analyzer (FIAstar 5000, FOSS, Hillerød, Denmark). The elemental composition (C and N) of bacteria was determined at the beginning and the end of the experiment. Water from each treatment was filtered onto a pre-combusted and pre-weighted Advantec GF-75 glass fiber filter (25 mm, 0.3µm pore size). The filters were pre-combusted at 450 °C in a furnace (Nabertherm LT 5/11/P33) for 5 hours. The filters were weighted to determine the bacterial dry mass produced during the experiment. The C and N content of bacteria was measured with a CHN elementary analyzer (Costech Elemental Combustion System 4010).

### Calculated variables

Based on the data we analyzed from water chemistry and bacterial stoichiometry, bacterial yield (*Y*_*C*_ and *Y*_*N*_) was calculated as the change in bacterial C or N content between the end and the beginning of the experiment:

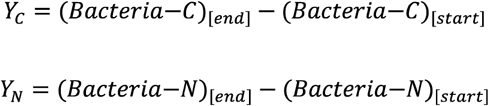

The C and N consumptions (*U*_*C*_ and *U*_*N*_) were calculated as the changes in substrate concentration between the end and the beginning of the experiment.

First, we estimated DON (dissolved organic nitrogen) as equal to TDN from which the dominant forms of inorganic nitrogen were subtracted:

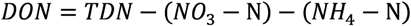

For C, we estimated consumption *U*_*C*_ as

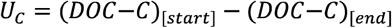

Consumption of N was estimated as the drawdown in the N-containing substrate(s) in each treatment. Hence, in the DIN/DOC treatments, where the nitrate was the source of N,

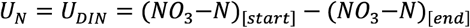

in the DON treatments, the N was provided entirely as DON, thus

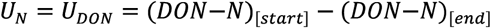

Last, in the COMBINE treatments, N was provided both as nitrate and as DON, thus

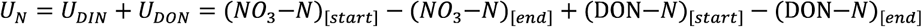

The bacterial ammonium regeneration (*M*_*N*_) was calculated as:

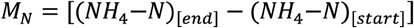

Net nitrogen mineralization (*NNM*) was calculated as the balance between ammonium regeneration and N immobilization:

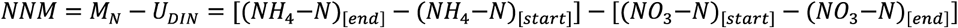

The bacteria growth efficiency for C and N (*BGE*_*C*_ and *BGE*_*N*_) was estimated as the proportion of the consumed C or N that was assimilated in new biomass.

For C, since we did not measure respiration rates, we used bacterial yield as a measure of net assimilation:

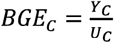

For N, we estimated net assimilation as N consumption, *U*_*N*_, minus N lost from bacteria to the medium as ammonium, *M*_*N*_:

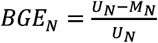

### Model predictions

Since C and N consumption (i.e., *U*_*C*_, *U*_*N*_, *U*_*DIN*_, *U*_*DON*_) and ammonium regeneration *M*_*N*_ can be directly calculated in our experiment, the C: N consumption ratio can be calculated independently from the supply ratio *S*_*C:N*_ or *O*_*C:N*_. Hence, we use the real consumption ratio to replace *S*_*C:N*_ (in equation 7), and *O*_*C:N*_ (in equation 9) in order to test the potential negative effect of assumptions (5) and (10) on the accuracy of ammonium regeneration predictions.

Accordingly, for equation (7) we calculated *M*_*N*_ a second time as:

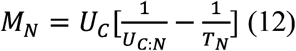

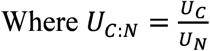

In equation (9) we recalculated *NNM* as

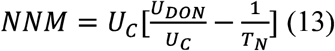

Finally, for the DIN/DOC treatment, nitrogen consumption in the model is assumed to be equal to 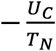 (see equation 11). In that last case, we replaced the assumed nitrogen consumption by its measured counterpart, yielding:

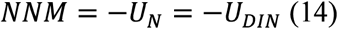

### Statistical analyses

All statistical analyses and figures were completed using R (3.5.1). Before statistical analysis on the stoichiometry data of bacteria community, we analyzed the bacteria abundance of the various treatments. As there was no bacteria growth in the alanine addition compared to the control treatment, part of the data we collected from the L-alanine treatment (e.g. bacteria yield, carbon and nitrogen consumption) were very close to or below the detection limit of instruments. Therefore, we removed all the data from the alanine treatment to avoid bias in statistical analyses. We conducted one-way ANOVAs to test for a significant treatment effect, using each treatment as a separate level. Differences between treatments were then tested using Tukey’s post-hoc test in ‘‘Agricola’’ package. All statistical tests used a family-wise significance level of 5% (*α* = 0.05).

## Results

At the end of our experiment, the bacterial abundance was very low in the control treatment, which was expected given the lack of any organic carbon substrate. In one of the DON treatments, where L-alanine was added, the bacterial abundance was similarly low (Fig. 1), which meant that most chemical analyses did not reach the detection limit. The data from this treatment are therefore not presented in the reminder of the result section.

**Figure 1.**
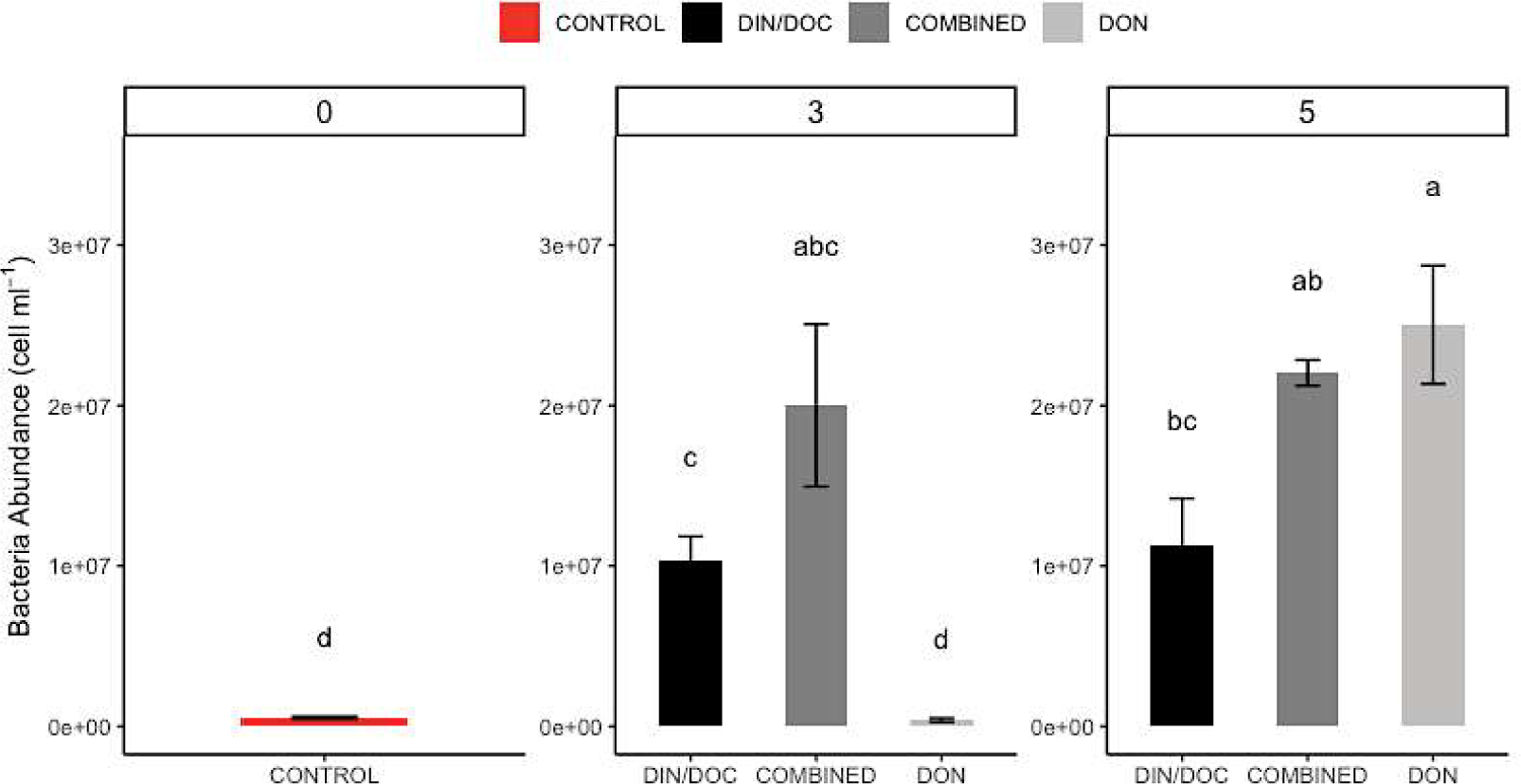
Bacteria abundance response to substrates of different molecular composition at the end of the experiment. The substrate supply C:N ratio (*S*_*C:N*_) (0,3 and 5) is indicated over the two groups. The mean ± se is plotted and different letters indicate statistical differences between treatments (Tukey’s HSD test, *p* < 0.05).

The other treatments besides DON-3 showed a significant increase in bacterial abundance at the end of the experiment, especially in the DON-5 treatment, in which C and N were associated in one molecular compound, glutamic acid. In both treatments where C and N were supplied in two separate molecular compounds, DIN/DOC-3 and DIN/DOC-5, bacterial abundance was around half the abundance in the DON-5 treatment on average (Fig. 1). In the COMBINED treatments where C and N were provided both associated in one molecule and dissociated in two molecules, the bacterial community reached mean abundance levels intermediate between the DON and the DIN/DOC treatments, although the difference with each of these treatments were not statistically significant (Fig. 1).

There was no ammonium regenerated (Fig. 2A), but significant nitrate uptake at the end of the experiment in the two DIN/DOC treatments (Fig. 2B). In the COMBINED treatments, bacteria did both take up nitrate and regenerate ammonium, resulting in a moderate net uptake of inorganic nitrogen (Fig. 2B).

**Figure 2.**
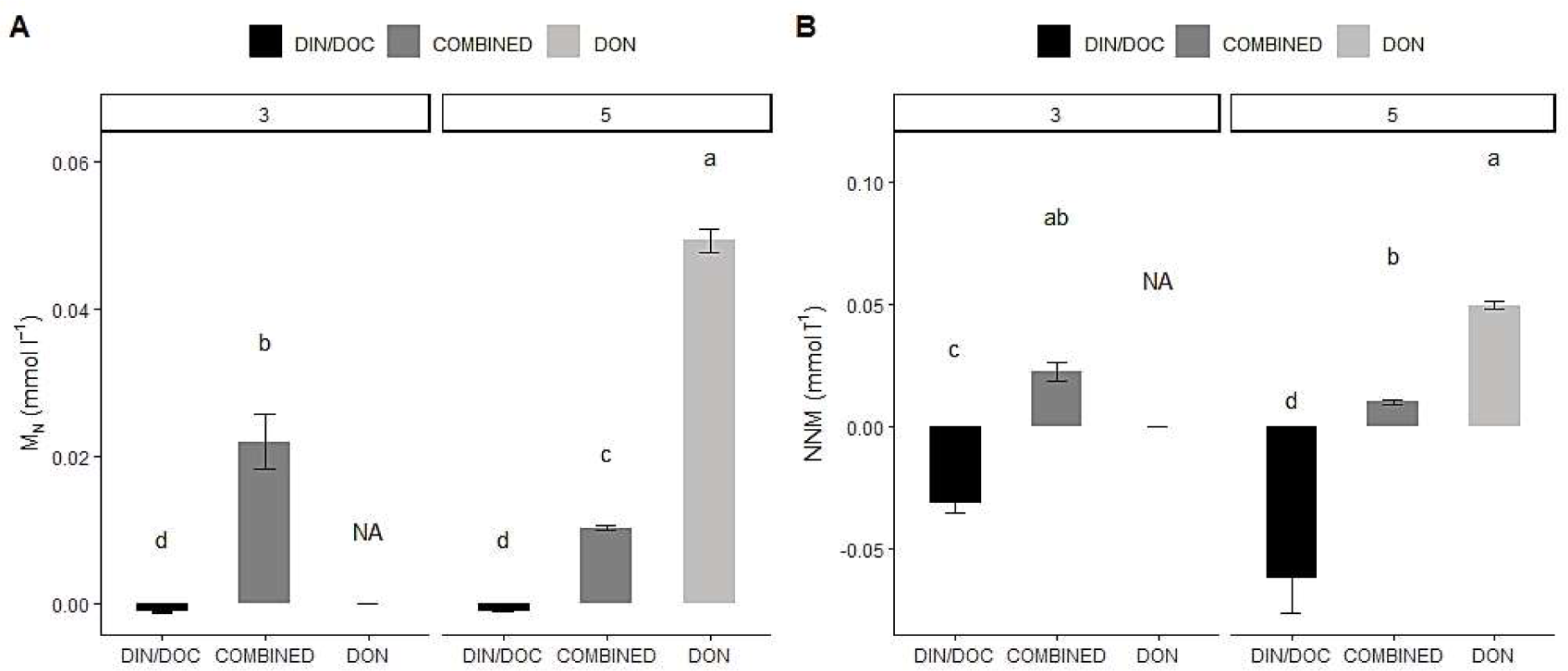
Effect of different substrates on A) ammonium regeneration (*M*_*N*_), and B) net nitrogen mineralization (*NNM*). The substrate supply C:N ratio (*S*_*C:N*_) (3 and 5) is indicated over the two groups. The mean ± se is plotted and different letters indicate statistical differences between treatments (Tukey’s HSD test, *p* < 0.05).

The bacterial yield C:N ratios (*Y*_*C:N*_) were relatively stable across treatments (Fig. 3C). However, the bacterial communities appeared to differ in their C and N yields across treatments (Fig. 3A, 3B). Overall, the DON-5 treatment yielded significantly more C and N biomass than other treatments.

**Figure 3.**
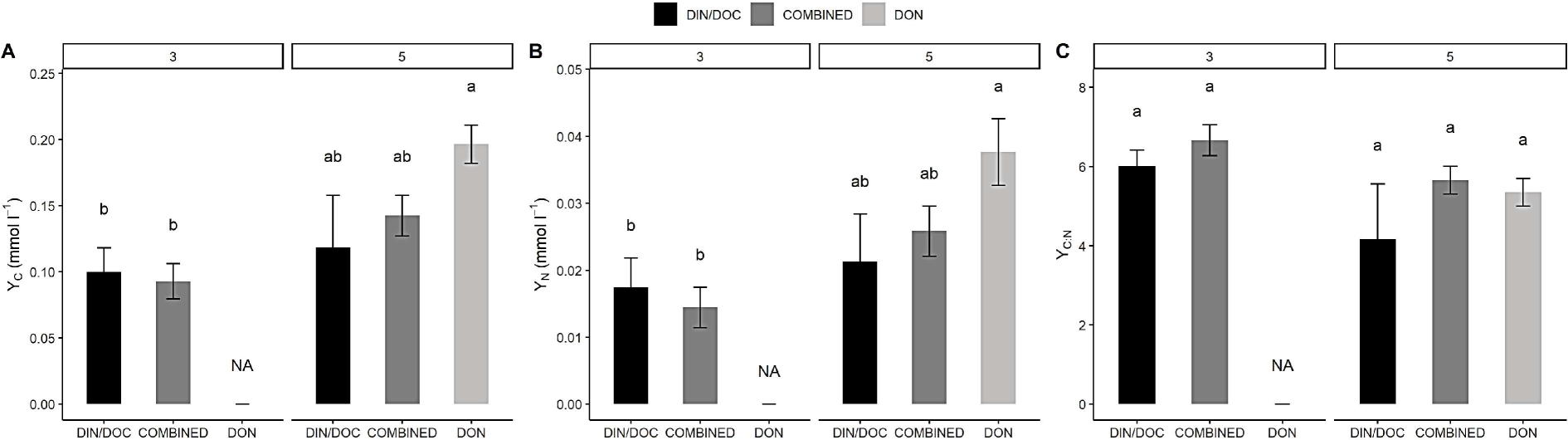
Effect of different substrates on A) bacterial carbon yield (*Y*_*C*_), B) bacterial nitrogen yield (*Y*_*N*_), C) bacterial C: N ratio in yield (*Y*_*C:N*_). The substrate supply C:N ratio (*S*_*C:N*_) (3 and 5) is indicated over the two groups in each graph. The mean ± se is plotted and different letters indicate statistical differences between treatments (Tukey’s HSD test, *p* < 0.05). NA: missing data.

Patterns in C and N consumption reflected those in C and N yields for treatments with a substrate supply ratio of 5 (Fig. 4). In the treatments with a supply ratio of 3, bacteria consumed significantly less C and N than the treatments with a supply ratio of 5 (Fig. 4A, 4B). Even though similar yields were observed among treatments with a supply ratio of 3 (Fig. 3), bacteria in DIN/DOC-3 consumed significantly less C and N than in the COMBINED-3 treatment. Despite the large differences in C and N consumption observed among most treatments, the ratio of C:N consumption was relatively stable across treatments, with only DIN/DOC-3 showing significantly lower consumption C:N ratios (Fig. 4C).

**Figure 4.**
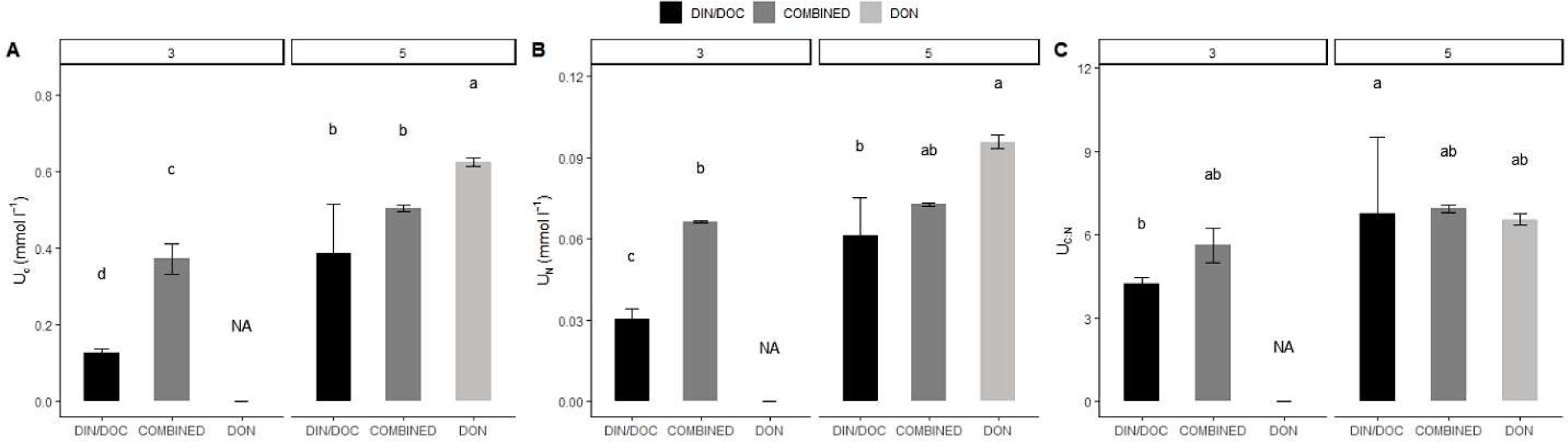
Effect of different substrates on A) bacterial carbon consumption (*U*_*C*_), B) bacterial nitrogen consumption (*U*_*N*_) and C) bacteria C: N consumption ratio (*U*_*C: N*_). The substrate supply C:N ratio (*S*_*C:N*_) (3 and 5) is indicated over the two groups in each graph. The mean ± se is plotted and different letters indicate statistical differences between treatments (Tukey’s HSD test, *p* < 0.05). NA: missing data.

The lower C consumption observed in the DIN/DOC-3 treatment was compensated by a higher bacterial carbon growth efficiency (Fig. 5A), thus explaining the similar C yields to the COMBINED-3 treatment. In all the other treatments, the bacterial community showed a relatively constant *BGE*_*C*_, at around a value of 0.25 (Fig. 5A). Differing from the consumption and yield, *BGE*_*N*_ showed clear differences between treatments (Fig. 5B) that mostly reflected the differences in ammonium regeneration shown in figure 1A. The DIN/DOC treatments, where no ammonium regeneration was found, achieved the highest *BGE*_*N*_, followed by the COMBINED treatments, where some ammonium was regenerated, and finally by the DON-5 treatment, whose bacteria regenerated the largest amounts of ammonium.

**Figure 5.**
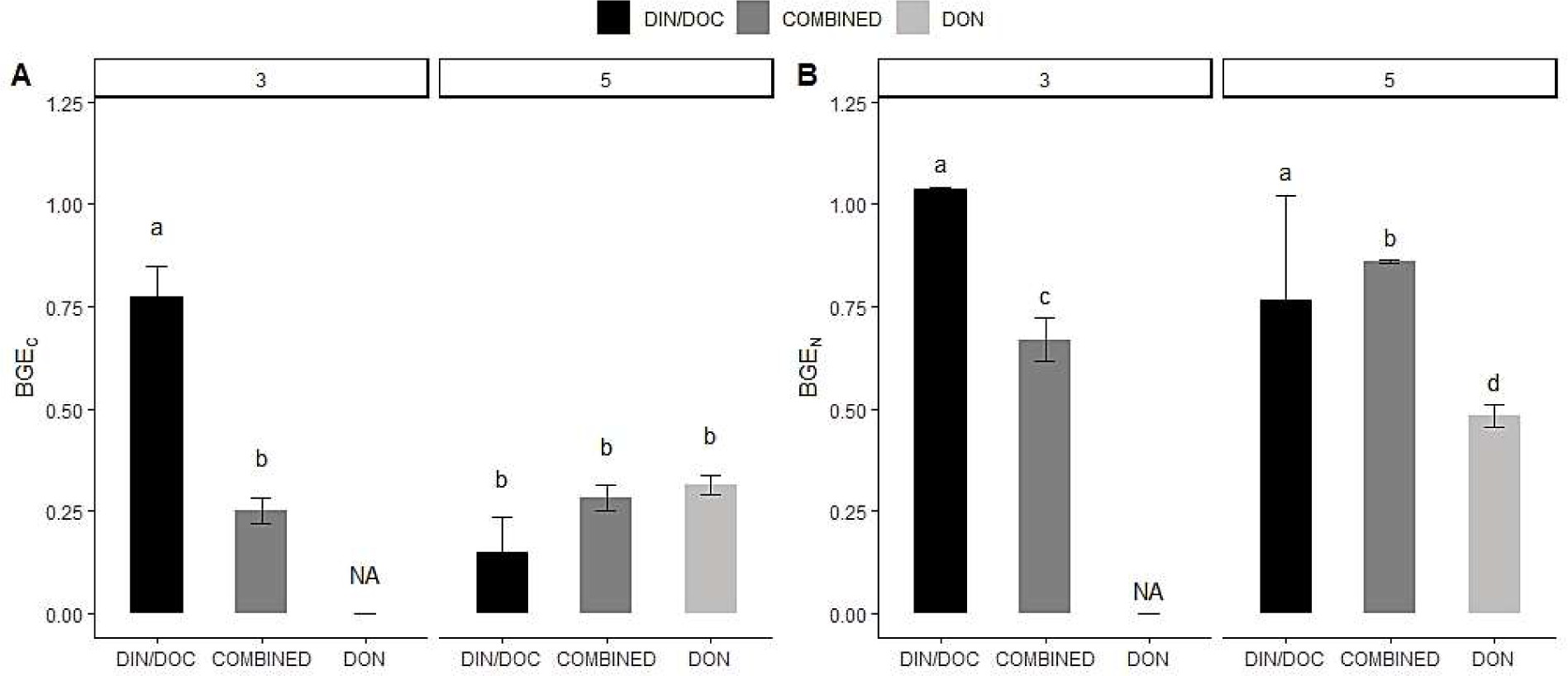
Effect of different substrates on A) bacterial carbon growth efficiency (*BGE*_*C*_) and B) bacterial nitrogen growth efficiency (*BGE*_*N*_). The substrate supply C:N ratio (*S*_*C:N*_) (3 and 5) is indicated over the two groups in each graph. The mean ± se is plotted and different letters indicate statistical differences between treatments (Tukey’s HSD test, *p* < 0.05). NA: missing data.

The simultaneous measurement of *Y*_*C:N*_ and *BGE*_*C*_ allows for the estimation of the net nitrogen mineralization threshold, *T*_*N*_, using equation (6). We found very different *T*_*N*_ even between treatments with similar supply ratios *S*_*C:N*_ (Fig. 6). In the DIN/DOC treatments, the average of *T*_*N*_ was close to 11:1. The other treatments had mean *T*_*N*_ around 22 with the exception of the COMBINED-3 treatment, whose mean threshold ratio was the highest (Fig. 6).

**Figure 6.**
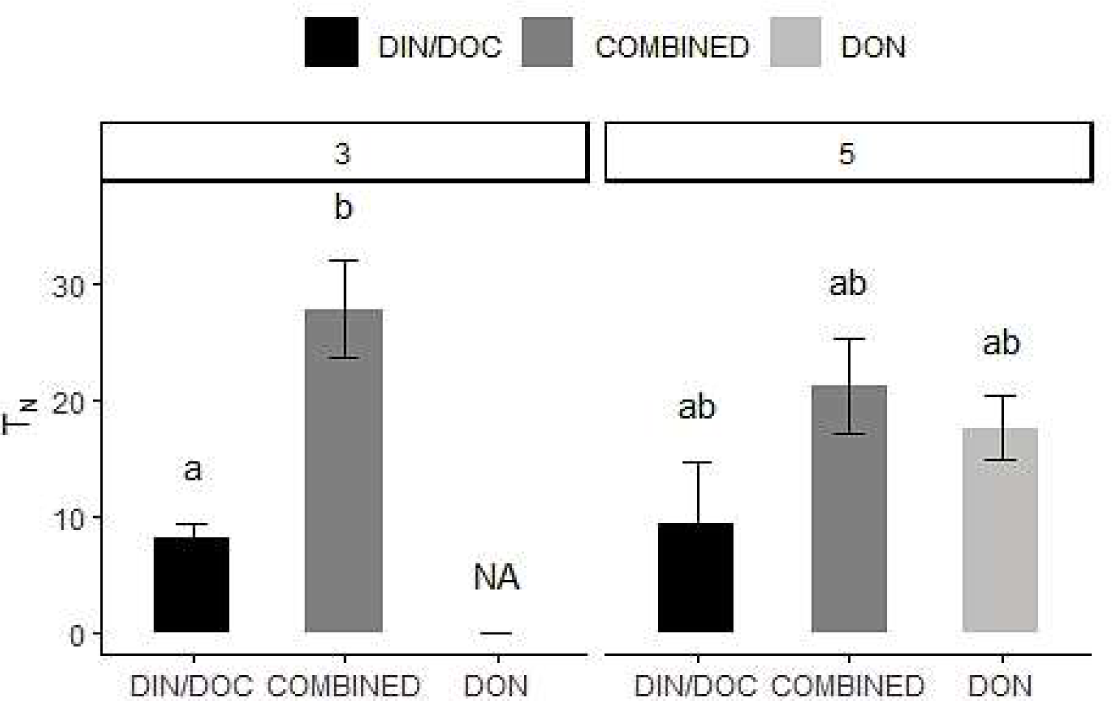
Effect of different substrates on the net nitrogen mineralization threshold (*T*_*N*_). The substrate supply C:N ratio (*S*_*C:N*_) (3 and 5) is indicated over the two groups. The mean ± se is plotted and different letters indicate statistical differences between treatments (Tukey’s HSD test, *p* < 0.05). NA: missing data.

Finally, we compared ammonium regeneration (*M*_*N*_) that we measured in our experiment with the predictions from equation (7) and equation (12) (Fig. 7A). The consumption C:N ratio (*U*_*C:N*_) was a better parameter to estimate *M*_*N*_ than the substrate C:N ratio (*S*_*C:N*_) in Goldman’s model. The predictions were clearly closer to measurements for the DON treatment, and for all treatments when *U*_*C:N*_ was used instead of *S*_*C:N*_ in equation (7). However, even using *U*_*C:N*_, the predicted *M*_*N*_ in the COMBINED treatments was still far from measurement. This suggests that for the dual N substrates, none of the estimates were good enough for predicting *M*_*N*_ accurately. On the other hand, the comparison between measured net mineralization (*NNM*) and the predictions by using equation (9) and (13) showed that the predictions using real consumption ratio were not always more accurate (Fig. 7B). Equation (13) correctly predicted *NNM* in both DIN/DOC and DON treatments, but not for the COMBINED treatment.

**Figure 7.**
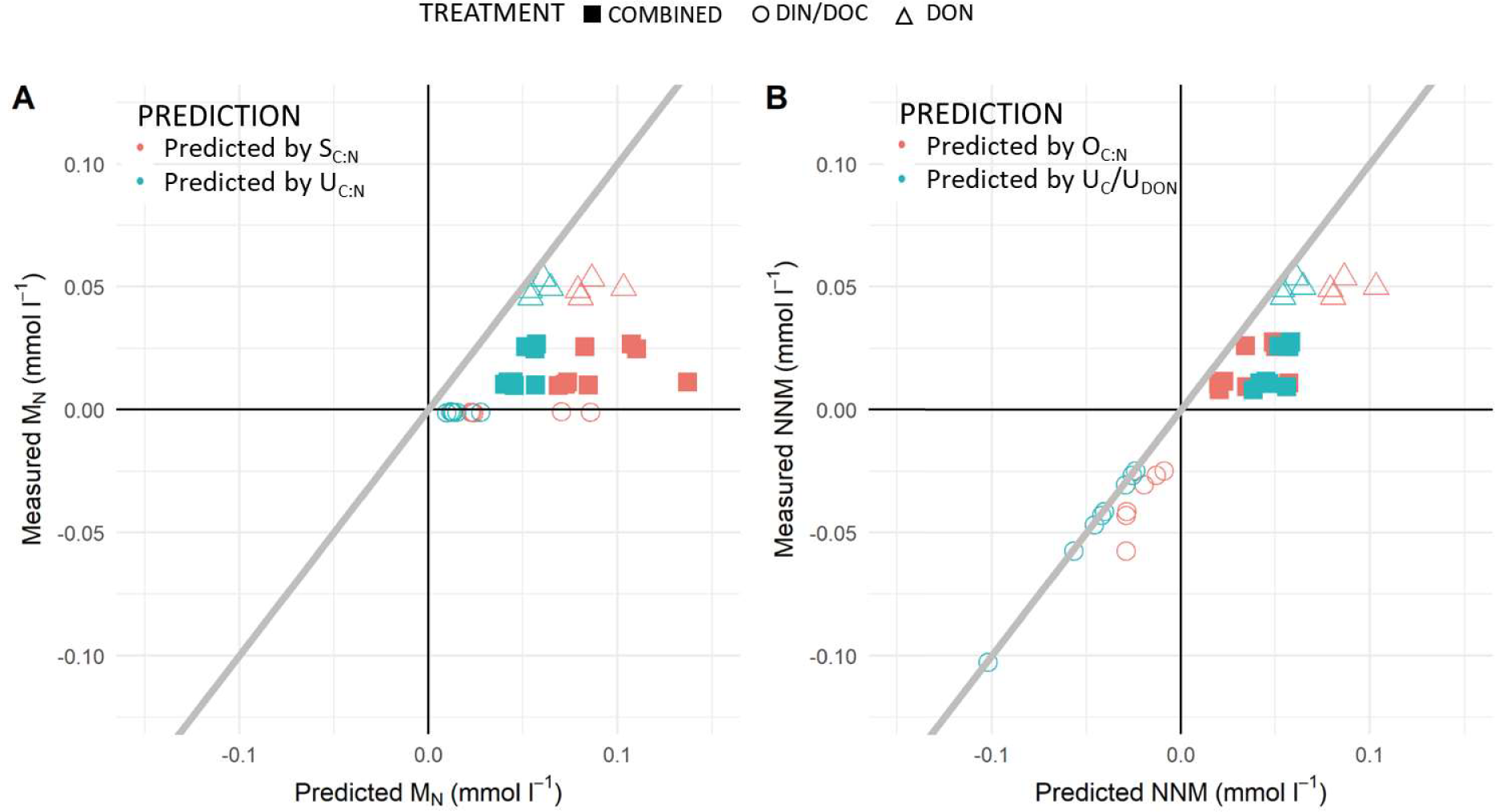
Measured A) ammonium regeneration (*M*_*N*_) and B) net nitrogen mineralization (*NNM*) for all treatments compared to the corresponding *M*_*N*_ and *NNM* predicted from equation (7) and equation (9) using substrate C:N ratio and measured real consumption C:N ratio as estimates of the consumed substrate C:N ratio. The closer a prediction is from the 1:1 grey line, the better it matches the real measurement.

## Discussion

Our experiment shows that, beyond elemental stoichiometry, the molecular composition of bacterial substrates plays a major role in both ammonium regeneration and net nitrogen mineralization. Growing on substrates with different molecular composition but with similar C:N ratios (*S*_*C:N*_), bacterial communities showed significantly different ammonium regeneration and net nitrogen mineralization. Therefore, we conclude that using *S*_*C:N*_ to predict bacterial *M*_*N*_ may in many cases not suffice. Actually, we found that using the stoichiometric consumption ratio, *U*_*C:N*_, instead of *S*_*C:N*_, improved the predictions significantly.

Of the different properties that define molecular composition beside stoichiometry, we focus on the level of association between C and N in the substrate molecules, as it is the property that we manipulated in our experiment. Indeed, in the DIN/DOC treatments, we had C and N fully dissociated, with substrate N contained in nitrate, and substrate C in a keto-acid (pyruvate or *α*-ketoglutarate). Because we selected substrates with low C:N ratios (3:1 and 5:1 respectively), the application of equation (7) to these treatments predicts significant ammonium regeneration in the DIN/DOC treatments. Our use of nitrate as the inorganic substrate, rather than ammonium (as in most previous similar experiments, e.g., Goldman *et al.* 1987; Goldman & Dennett 1991, 2000), allowed us to detect that there were no or very little ammonium regeneration in these treatments (Fig. 1A and 7A). Thus, our results suggest that bacteria balanced their relative assimilation of C and N to their needs entirely by adjusting their C and N consumption and not through the excretion of ammonium. Hence, in cases where N and C are fully dissociated, we expect no ammonium regeneration, unlike what equation (7) predicts.

Secondly, in the COMBINED treatments, we manipulated the C and N association in a different way, using a mixture of inorganic (N-only substrates) and organic substrates (both C-only and C- and-N-containing substrates). In these treatments, bacteria may actively choose different form of substrates that they need to balance their uptake of C and N, and thus, consumption C: N ratio do not fully reflect the substrate C: N ratio, explaining why equation (7) works better when using *U*_*C:N*_ instead of *S*_*C:N*_ (Fig. 7). However, because a substantial fraction of C is linked to N in amino acids, bacteria do not take up N independently from C, and thus need to excrete the excess of N taken up as ammonium (Fig. 1).

When it comes to DON treatments, results were, at first sight, unexpected. First, bacteria did not grow in any substantial way in the DON-3 treatment. In this treatment, L-alanine was used as the C and N substrates. Hence, we are drawn to conclude that our bacterial community could not grow on L-alanine as its only organic substrate, although previous researchers have grown bacterial strains on this substrate (Franklin & Venables 1976; Goldman *et al.* 1987). It seems that high concentrations of L-alanine may be toxic for some bacteria as found by Kim et al (2015), while other bacteria have the necessary enzymes to use the amino-acid for growth (Coudert 1975). An alternative explanation could be that growing on amino acids requires deamination in order to use the carbon backbone for respiration or further cell building. Among the enzymes involved in deamination are the amino-acid oxidases, which produce hydrogen peroxide in stoichiometric amounts to the amino-acids oxidized (Hossain *et al.* 2014). Hydrogen peroxide is known to be highly toxic to bacterial cells without catalase (the enzyme that detoxifies hydrogen peroxide), to the point that some amino-acid oxidases can be used as anti-microbial compounds (Hossain *et al.* 2014). One hypothesis we put forward is that, because alanine has a very low C:N ratio (3:1), bacteria need to deaminate a substantial proportion of the alanine they assimilate (at least 80% if we assume a *BGE*_*C*_ of 20%). This process should generate a lot of hydrogen peroxide (at least one molecule for every 3 C atom respired) and thus be toxic to most, but not all, bacterial strains (Goldman *et al.* 1987). In the DON-5 treatment, our bacterial community managed to grow well on glutamate, an amino acid with a C:N ratio of 5. Since C and N were completely associated in the amino-acid, one would expect *U*_*C:N*_ to be equal to *S*_*C:N*_ in this treatment. But our results showed that equation (12) – when *U*_*C:N*_ was used – still performed better than equation (7) that used *S*_*C:N*_ (Fig. 7). Again, we hypothesize that the deamination process is critical to understand this result. Some of the bacterial L-Amino acid oxidases needed for this process are extracellular (Hossain *et al.* 2014). Thus, it is likely that a fraction of the C and N in this treatment was dissociated already outside of the cells, allowing for the partial adjustment of *U*_*C:N*_ to the needs of the bacteria. Overall, in the COMBINED and DON treatments, the consumption C:N ratio, *U*_*C:N*_, was a better predictor of M_N_ than *S*_*C:N*_, which likely reflects that the partial dissociation of C and N in the substrate allows for an adjustment of relative C and N consumption.

We measured other parameters in equation (7) besides *U*_*C:N*_, which are important in determining ammonium regeneration. *T*_*N*,_ the stoichiometric net mineralization threshold, marks the limit between the *U*_*C:N*_-values that result in net immobilization from those that result in net mineralization. As described in equation (6), *T*_*N*_ is determined by both *Y*_*C:N*_, the bacterial biomass C:N ratio, and *BGE*_*C*_, the bacterial growth efficiency for C. In our experiment, we found that *Y*_*C:N*_ was relatively stable among treatments as was found in many studies in which bacteria were grown on substrates with low C:N ratios (Mooshammer *et al.* 2014). Other studies, however, found that bacteria had more variable biomass C:N ratios when grown on substrates with high C:N ratios (Stenzel *et al.* 2017). By using only two substrate C:N ratios (3 and 5) our experiment might thus have missed some regulatory mechanisms besides the adjustment of relative C and N consumptions, which may affect *M*_*N*_ under conditions of high C:N supply ratios. *BGE*_*C*_ was also relatively constant across treatments. The observed values were around a value of 0.25, which is typical of bacteria growing under good conditions (Del Giorgio & Cole 1998), with one notable exception: bacteria growing in the DIN/DOC-3 treatment, i.e., growing on pyruvate, showed elevated efficiency, around 0.75. This is no surprise, given the central role of pyruvate in bacterial metabolism (Cook 1930). Also, when C and N are not associated in one molecule, such as in DIN/DOC-3, the degree of freedom in adjusting the relative uptake of C and N decreases the chance of taking up C in excess, which increases the carbon growth efficiency (Diner *et al.* 2016). Hence, *BGE*_*C*_, and by extension *T*_*N*_, depends like M_N_ on the molecular composition of the substrate and on the degree of association between N and C, via the possibility for bacteria to adjust their relative consumption of C and N.

## Conclusion

The purpose of this study was in part to test a widely used stoichiometric model of ammonium regeneration, by varying the molecular composition of the supplied substrates, and to propose its revision so as to reach better predictions. We found that, whenever possible, measuring the actual consumption C: N ratio is crucial. Natural aquatic ecosystems are likely to show even higher degrees of substrate heterogeneity than in our experiment. Hence it is all the more important to try to estimate the degree of association between C and N in natural systems of interest. It is still very difficult to characterize DOM composition precisely, but Fourier Transform Ion Cyclotron Resonance Mass Spectrometry (FT-ICR MS) or Orbitrap is a promising method that provide information on C bonds to other elements such as O, H and N (Koch *et al.* 2005; Simon *et al.* 2018). Conceptually, the possibility for bacteria to fulfill their stoichiometric needs by adjusting their relative consumption of C and N means that bacteria are less likely to affect the availability of inorganic N via ammonium regeneration than is predicted by models which base their estimates on equation (7). Hence, the role of bacteria in indirectly controlling the growth of phytoplankton by shifting from immobilization to mineralization as a function of their substrate C:N ratio, and thus switching the bacteria-phytoplankton interaction from competition to mutualism, might be less important than assumed.

## Acknowledgements

We acknowledge the financial support of the Knut and Alice Wallenberg Foundation via the project “Climate change induced regime shifts in Northern lake ecosystems”. We thank Anders Jonsson from Biogeochemical analytical facility KBC infrastructure for chemical analyses and Sonia Brugel, Renyuan Huang for their assistance in the lab. We also thank Sebastian Diehl, Ryan Sponseller and Göran Englund for their valuable comments on this manuscript.

